# NMR Spectral Alignment Utilizing a CryoEM Motion Correction Algorithm

**DOI:** 10.1101/2025.05.05.652277

**Authors:** Colin A. Hemme, Owen A. Warmuth, Songlin Wang, Christopher G. Williams, Alexander Thome, Leonard J. Mueller, Chad M. Rienstra, Timothy Grant

## Abstract

With recent advances in magic-angle spinning (MAS) solid-state NMR (SSNMR) resolution, precise spectral alignment has become a critical bottleneck in data processing workflows. While solution NMR employs deuterium lock systems, most SSNMR probes still lack this capability; though a lock corrects for magnet drift and instabilities, it is not alone sufficient to account for field gradients, sample temperature differences, and pulse sequence effects that can contribute to referencing errors among several data sets. These offsets become particularly problematic in the lengthy multidimensional experiments that provide the foundation for resonance assignment and structure determination procedures. Currently, researchers rely on manual alignment through visual peak inspection—a qualitative approach that often overemphasizes prominent, outlying peaks while overlooking subtle, global patterns. This subjective process becomes increasingly impractical for use cases with lower sensitivity, such as large proteins with thousands of peaks. To address these challenges, here we present Automated NMR Spectral Alignment (*ANSA*), a program that adapts cryo-electron microscopy motion correction principles to NMR spectroscopy. *ANSA* treats NMR spectra as images and applies cross-correlation functions to determine optimal alignment, improving cross-correlation scores from 0.33 to 1.00 in controlled tests and achieving 0.96 correlation in real-world applications with previously misaligned spectra. The algorithm successfully aligns spectra across varying experimental conditions, corrects shifts in long-duration experiments, and works with 2D and 3D datasets, with approaches that can be readily extended to additional dimensions. By eliminating human bias and providing objective, consistent spectral alignment, *ANSA* enhances scientific rigor, improves reproducibility between experiments, and enables automation of critical data processing steps. The software is freely available as an open-source tool, ready for integration into existing NMR workflows.

## INTRODUCTION

Nuclear magnetic resonance (NMR) remains a cornerstone technique for investigating systems from small organic molecules to large biomolecular assemblies^1,2,3,4^. As a high-precision and powerful tool in structural biology, NMR enables atomic-resolution structure determination and detects subtle structural perturbations through the exquisite sensitivity of chemical shifts to even minor changes in a local electronic environment. With the advent of ultra-high field magnets (>1 GHz Larmor frequencies), multidimensional pulse sequences, improved decoupling schemes^5–8^, and advanced probes^9–11^, SSNMR resolution has improved dramatically, allowing investigation of increasingly complex systems. However, this enhanced resolution demands correspondingly precise spectral alignments, since even small referencing discrepancies can lead to misinterpretation of data^12^. The data analysis challenge scales with the molecular weight of the biological system, limiting routine NMR structure determination to ∼50 kDa^12^.

Meanwhile, single-particle cryo-electron microscopy (cryoEM) has over the last decade gained a significant increase in popularity owing to increases in achievable resolution, now routinely achieving sub-3.5 Å resolution maps that enable atomic modeling^13^. This “resolution revolution” has largely been fueled by direct electron detectors with superior efficiency^14^, transforming cryoEM data collection from single static images to movies and enabling correction of sample movement during beam exposure^15^—a critical preprocessing step for achieving high resolution.

Recently, the complementary strengths of NMR and cryoEM have spurred the development of integrated structural biology workflows, with hybrid approaches successfully characterizing symmetric assembles where the asymmetric subunit is amenable to NMR analysis and the larger assembly is amenable to cryoEM analysis^16^. Other implementations use cryoEM density as constraints in NMR structure calculations, combining cryoEM with high-precision NMR refinement^17^. These integrative results highlight a fundamental similarity between the techniques: both rely on averaging numerous independent signals to generate their output.

This signal averaging principle demands that data be precisely superimposable; independently collected datasets must align *perfectly*, i.e., within the digitization limits of the highest resolution data collected independently to be added together such that the resultant signal-to-noise ratio is increased. In cryoEM, this is achieved by various image processing steps, for example, when aligning raw movie data, or when aligning individual particle images^15,18^. In NMR, this critical alignment function has traditionally been performed through spectral referencing^19^—a process that, unlike its cryoEM counterpart, has resisted comprehensive automation and remains largely dependent on subjective human intervention. Even in solution NMR of proteins, where deuterium lock systems have long been available^20^ and standard referencing procedures agreed upon^21^, the temperature dependence of lock solvent chemical shifts often causes discrepancies that are handled through linear correlation analysis of a subset of the peaks^22^. This discrepancy presents an opportunity to adapt proven cryoEM alignment methodologies to address a persistent challenge in NMR data processing.

In this paper, we present Automated NMR Spectral Alignment (*ANSA*), an algorithm that adapts cryoEM motion correction principles and image processing techniques to NMR spectroscopy. In cryoEM, as the electron beam contacts the sample, it causes the specimen to move in a phenomenon known as beam-induced motion^15^, requiring alignment of individual frames of the movie. This alignment procedure, therefore, must be fully automatic and robust even at the very low signal-to-noise (SNR) ratios and must leverage the full image, utilizing weighted cross-correlation functions (CCF) between individual frames and the sum of all other frames to estimate shifts for that frame. The process is repeated iteratively, keeping shifts updated from the previous round until convergence^18,23^.

With *ANSA*, we consider NMR spectra as continuous images rather than discrete peaks, enabling us to essentially align spectra as though they were cryoEM movie frames. Using NMR spectra of a sample under different data collection conditions (different magnetic field, pulse sequence, temperature, spinning rate, etc.), a global cross-correlation function will pinpoint the global referencing value at which the spectra are most similar. Crucially, the approach is robust to noise and requires only a subset of the spectral dimensions and cross peaks to be common among the spectra.

*ANSA* is readily applicable to several common use cases. First, we show the accurate alignment of 2D spectra of controlled, artificial offsets with perfect accuracy. Second, we apply *ANSA* to correct for referencing errors among replicate spectra, with nominally the same conditions but collected with different spectrometer calibrations, months apart. Third, we show that *ANSA* can accurately align 2D ^13^C-^13^C correlation spectra at varied mixing times: datasets that may contain the same peaks but may also exhibit additional correlations at higher mixing times. Finally, we extend the approach to handle the complex case of aligning 3D spectra with 2D projections, a frequent requirement in biomolecular NMR studies. In all of these applications, we demonstrate that *ANSA* provides objective, reproducible, and automatic alignment that eliminates human bias while improving the efficiency and accuracy in NMR data processing workflows.

## EXPERIMENTAL SECTION

### Sample Preparation of Toho beta-Lactamase

Uniformly ^13^C-^15^N Toho-1 was expressed and purified following previously published procedures (Tomanicek 2011)^24^. Protein microcrystals were grown by mixing the protein at 300-400 μM in 20 mM MES buffer at pH 6.5 with a crystallization buffer containing 7 mM spermine and 30% PEG-8k at a 1:1 ratio. Crystals were allowed to grow over 3-5 days at 4 °C. The sample was packed in a 1.6 mm Varian-style SSNMR rotor using customized sample packing devices (Olson 2024)^25^.

### SSNMR Spectroscopy of Toho β-Lactamase

NMR experiments were performed on a Bruker NEO 1.1 GHz spectrometer at NMRFAM using a Black Fox 1.6 mm HCN triple-resonance probe (Black Fox LLC) featuring 2H lock functionality. All spectra were collected at a 25 kHz MAS rate, and the VT temperature was set to -5 ºC. All spectra were referenced to DSS, using adamantane as a secondary external standard. The downfield ^13^C signal of adamantane was referenced to 40.48 ppm for this data and all others.

The 2D NCO was performed with the ^13^C carrier frequency set to 175 ppm and the ^15^N carrier set to 117.353 ppm. Data were collected with a spectral width of 100 kHz in the direct dimension with 3072 complex points and 25 kHz in the indirect dimension with 1024 complex points and 16 scans per FID. Long-Observation-Window Band Selective Homonuclear Decoupling (LOW-BASHD)^8^ was implemented in the direct dimension to decouple the Cα-C’ J-coupling using *τ*Dec=3.2 ms and 72.5 μs Gaussian π-pulses.

### Sample Preparation of Alpha-Synuclein Fibrils

Uniformly ^13^C-^15^N labeled ASyn monomer was expressed and purified as described previously^26^. Fibrils were prepared according to Bagchi *et al*. (2013)^27^ For isotopically labeled SSNMR samples, fibril seeds were generated according to the above protocol. Then, uniformly labeled monomer was added to the suspension of fibril seeds, and amplification was conducted in 50 mM Tris-HCl buffer and 100 mM NaCl, pH 7.6.

### SSNMR Spectroscopy of Alpha-Synuclein Fibrils

Magic-angle spinning (MAS) SSNMR experiments were conducted at 17.6 T (750 MHz 1H frequency) using a Varian NMR (Walnut Creek, CA) VNMRS spectrometer. The VT gas was set to a temperature of 0 ºC. The 17.6 T magnet was equipped with a Balun 3.2 mm probe. All spectra were referenced using adamantane as a secondary external standard.

The 2D carbon-correlation spectra were performed using dipolar-assisted rotational resonance (DARR)^28^ mixing between the indirect and direct dimensions with a 50 ms mixing time. They were collected with the ^13^C carrier frequency set to 97.438 in both dimensions. Data were collected with a spectral width of 100 kHz in the direct dimension with 2000 complex points and 50 kHz in the indirect dimension with 640 points and 8 scans per FID.

### Sample Preparation of Tryptophan Synthase

Uniformly labeled ^13^C-^15^N Salmonella typhimurium Tryptophan synthase in E. coli was expressed and purified as previously described^29^. Microcrystals were collected and washed with 50 mM Cs-bicine, pH 7.8, containing 8% PEG-8000, 1.8 mM spermine, and 3 mM N-(4’-trifluoromethoxybenzenesulfonyl)-2-aminoethyl phosphate (F9; a high affinity alpha site ligand) as previously described^30^.

### SSNMR Spectroscopy of Tryptophan Synthase

NMR experiments were performed on a Bruker NEO 1.1 GHz spectrometer at NMRFAM using a Black Fox 1.6 mm HCN triple-resonance probe (Black Fox LLC) featuring 2H lock functionality. All spectra were collected at a 25 kHz MAS rate, and the VT temperature was set to -5 ºC. All spectra were referenced to DSS, using adamantane as a secondary external standard.

The 2D carbon correlation experiments were performed using Combined 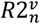 -Driven (CORD)^31^ mixing between the indirect and direct dimensions. For carbon-correlation experiments with 100 and 300 ms CORD mixing, the carrier frequency was set to 100 ppm. Any adjustment made to the carrier to highlight the functionality of *ANSA* was made relative to these values and is explicitly stated in the text. All 2D CC spectra were collected with a spectral width of 100 kHz and with 2048 complex points in the direct dimension and 800 complex points in the indirect dimension, with 8 scans per FID. The 2D NCA experiment was performed with the ^13^C carrier frequency set to 55.65 and the ^15^N carrier set to 118 ppm. Data were collected with a spectral width of 100 kHz in the direct dimension with 3072 complex points and 25 kHz in the indirect dimension with 640 complex points and 16 scans per FID. The NCACO 3D used to generate NCA projections was performed with the ^13^C carrier set to 55 ppm and the ^15^N carrier set to 117.45 ppm. The data are comprised of 3072 complex points in the direct dimension with a spectral width of 100 kHz, 80 complex points in the ^13^Cα dimension with a spectral width of 12.5 kHz, and 128 complex points in the ^15^N dimension with a spectral width of 8.333 kHz. Long-Observation-Window Band Selective Homonuclear Decoupling (LOW-BASHD)^8^ was implemented in the direct dimension to decouple the Cα-C’ J-coupling using *τ*Dec=3.2 ms and 72.5 μs Gaussian π-pulses. The spectrum was acquired using a 25% NUS schedule.

### Alignment Procedure

*ANSA* uses cisTEM^32^ as a code base in C/C++ for development. *ANSA* takes data in the form of two ft2 files, which are converted to text and header files using NMRPipe^33^. These files are provided as input to the *ANSA* program for coordinates and conversions. A binary file is available for download from the NMRFAM website (https://nmrfam.wisc.edu/software/), and the source code can be downloaded from the *cis*TEM GitHub repository: https://github.com/timothygrant80/cisTEM

## THEORY

To align the spectra correctly and output shifts in ppm, the spectral sampling (SS) and offsets due to differing origins need to be accounted for. Calculation of the spectral sampling is performed as shown in (eq. 1) using the spectral width (sw) and transmitter frequency (obs) in Hz to determine the size of the spectral image and convert pixel size into ppm for both the *x* and *y* dimensions. *n* denotes the size in pixels of the spectra in each respective dimension.

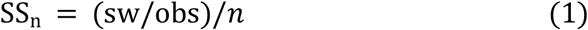

If the spectra have different origin values, the origin offset (O_offset_) must be calculated to output the correct ppm shift value. NMRPipe’s origin denotes the minimum ppm values of the spectra (i.e., the bottom-left) rather than the center values. This origin will therefore change as spectra are resampled and resized in preparation for alignment. Thus, to provide a general offset, we reframe the origin as the center of the spectral image. The O_offset_ (center) calculation is shown (eq. 2) using the dimensions of each spectrum (*n*) and the shared SS after resampling.

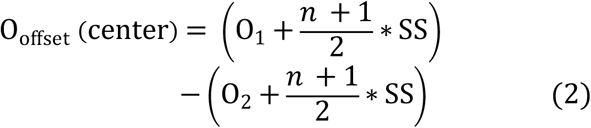

In cases where the spectra have different SS values, we Fourier interpolate the SS of spectrum 2 to spectrum 1. The calculation of the new resampled dimension for spectrum 2 is shown (eq. 3), where the spectra are resized to the new dimension in Fourier space.

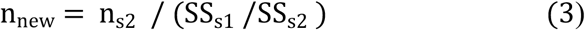

If spectra do not have the same dimensions, they are resized to the same physical dimensions while maintaining their center position. For 3D spectra, projections are generated using TCL scripts distributed with NMRPipe. To preserve SNR in the projections, the maximum-value mode is used by default. The resultant files can be treated identically to 2D datasets in *ANSA*. To reduce the effects of low-resolution features in the spectra (e.g., large density changes or very bright peaks), the spectra are high-pass filtered by Fourier transforming them, multiplying by an inverse Gaussian with a full width at half maximum of 0.2355, and then reverse Fourier transforming them. The spectra values are then zero-floated (set to an average value of 0) and normalized such that their variance is 1, ensuring that subsequently calculated cross-correlations will range between - 1 and 1. CCF is then calculated between the filtered, normalized spectra (eq. 4), with the peak of that function corresponding to the x,y pixel shifts that lead to maximum agreement between them. To provide finer “sub-pixel” shifts, a parabola function is fit to the CCF peak, and this parabola function is used to interpolate the shifts with sub-pixel resolution, increasing the accuracy of the determined shifts.

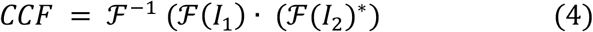

## RESULTS AND DISCUSSION

To validate *ANSA*’s alignment capabilities, we conducted a systematic series of tests progressing from controlled artificial offsets to increasingly complex real-world scenarios involving different experimental conditions. First, we tested *ANSA*’s ability to detect known chemical shift differences by artificially inducing a 0.1 ppm offset in a Toho β-lactamase NCO spectrum. Visual inspection confirmed the offset between the original and shifted spectra (Figure 1A), which was reflected in their low initial cross-correlation score of 0.33. *ANSA* precisely calculated the offset as 0.1 ppm in both dimensions—matching the known applied shift with ±0.01 accuracy. As expected for identical spectra differing only in their offset, alignment improved the cross-correlation score to 1.00 (Figure 1B), confirming that *ANSA* can quantitatively detect and correct chemical shift offsets with high precision. Next, we evaluated *ANSA*’s performance with real experimental data exhibiting natural referencing variations. We applied the algorithm to two Alpha-Synuclein Fibril spectra collected several months apart (June and November), representing a common scenario where spectrometer drift or different calibration settings create alignment challenges. In this case, a malfunction of the spectrometer shim power supply in mid-experiment caused an especially large error that would normally deem the data to be unusable. *ANSA* detected a substantial offset of (2.682, 2.692) ppm between these datasets. After applying this correction, the spectra over--laid remarkably well (Figure 2B compared to unaligned spectra in Figure 2A), with the cross-correlation score improving from 0.46 to 0.96. The slight deviation from perfect correlation (1.00) is expected and appropriate given the different noise profiles in these independently collected datasets. This result demonstrates *ANSA*’s ability to correct significant referencing discrepancies in real experimental data collected under varying conditions over extended periods—a critical capability for longitudinal studies and multiuser facilities.

**Figure 1.**
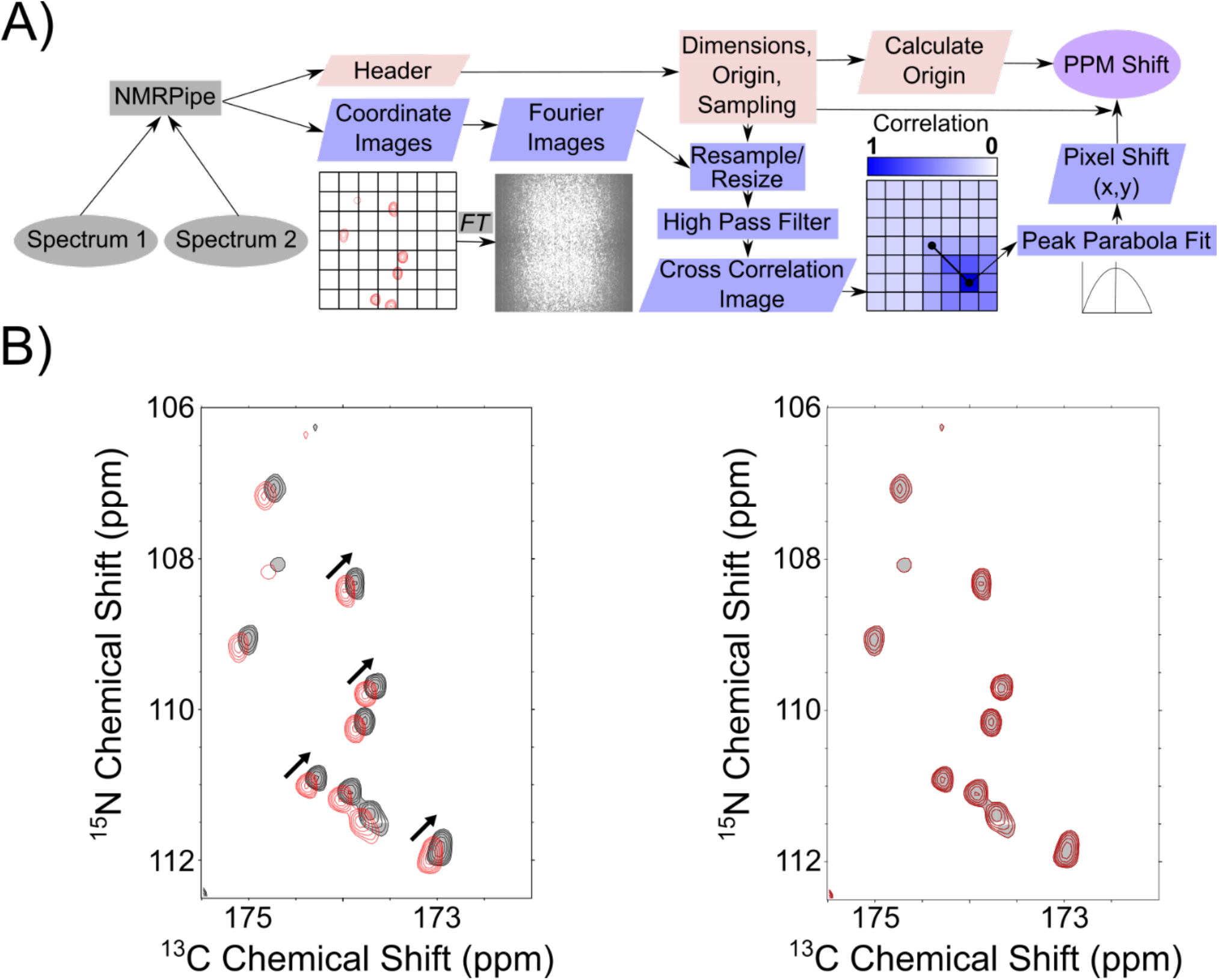
Schematic of *ANSA* and alignment of intentionally offset spectra. (A) Schematic of *ANSA* program workflow. *ANSA* takes two spectra as input, which are converted to images and header information. The images are appropriately resampled, resized, normalized, and filtered, then compared via CCF to determine the shift in pixels that leads to the maximum agreement between them. The header information of the spectra is parsed into the program, which is used to accurately convert the final pixel shifts into ppm values. (B) Overlaid NCO spectra of Toho1 β-lactamase with (red) and without (black) an applied offset of 0.1 ppm. The inlay highlights this offset. On the right are the spectra aligned with *ANSA*. The calculated offset was exactly 0.1 ppm in the ^15^N and ^13^C dimensions, highlighted by the exact overlap of peaks in the inlay. The cross-correlation score increases from 0.33 before alignment to 1.00 after.

**Figure 2.**
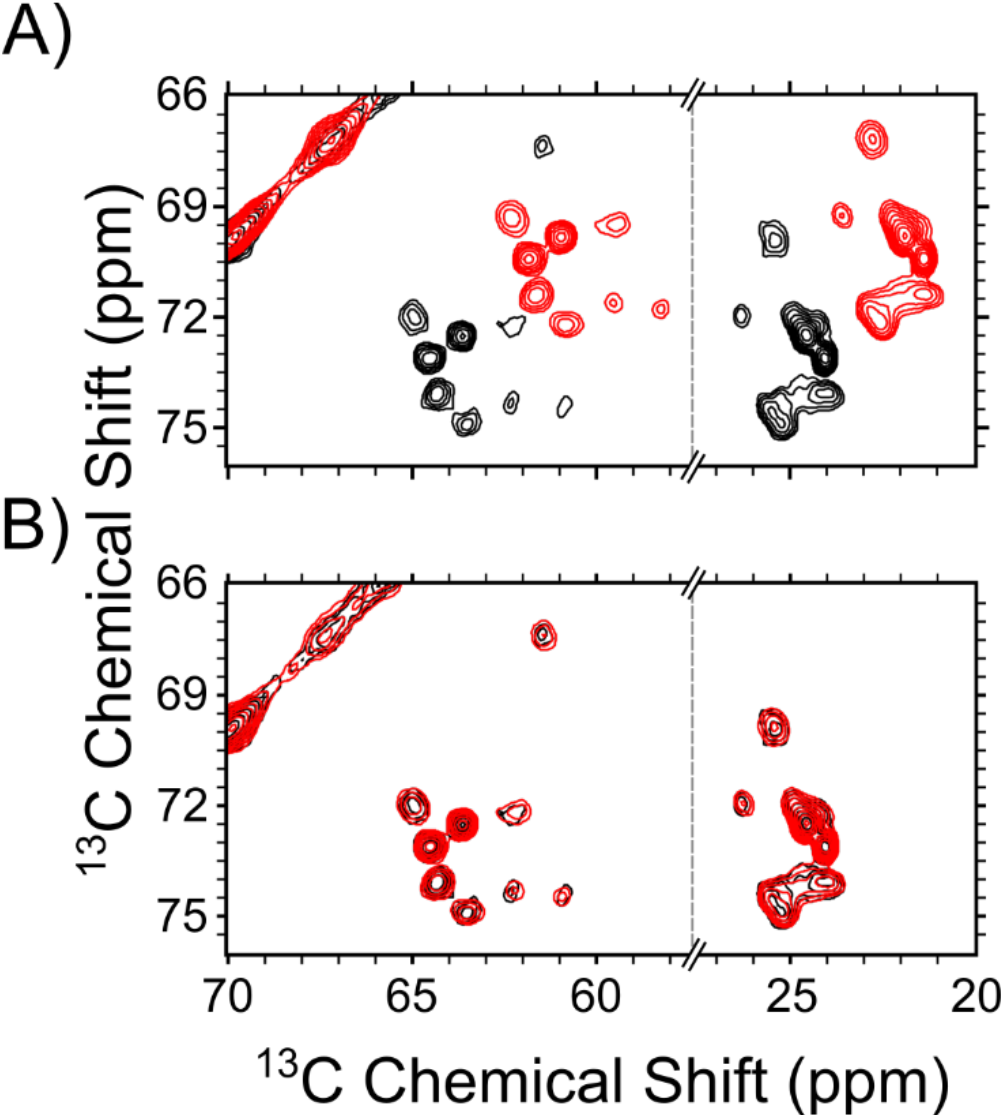
Alignment of experimental replicates collected at separate times. (A) Overlay of two unaligned Alpha Syn carbon-correlation spectra collected in June (black) and November (red). (B) Overlay of aligned spectra from (A). The detected shift was (2.682, 2.692) ppm, and the cross-correlation score was 0.46 before alignment and 0.96 after alignment.

To further challenge the algorithm, we examined alignment between spectra containing both shared and unique peaks. We applied *ANSA* to two tryptophan synthase ^13^C-^13^C correlation spectra collected with different CORD mixing times (100 ms and 300 ms), which contained both common peaks and additional correlations unique to the longer mixing time experiment. Despite these spectral differences and an obvious referencing error, *ANSA* successfully detected an offset of (0.580, 0.609) ppm. Application of this correction resulted in excellent visual alignment (Figure 3C) and improved the cross-correlation score from 0.83 to 0.93. To test the sensitivity of the alignment, we also applied a shift corresponding to the average of the detected values, (0.595, 0.595) ppm, and observed a slightly lower cross-correlation score of 0.92 which shows that an average shift is close but non-optimal. This result highlights that even though the x and y components of the optimal shift differ, ANSA accurately identifies the true best alignment. This result is particularly significant as it demonstrates *ANSA*’s robustness against variations in peak patterns and noise profiles—a key advantage over peak-picking based alignment methods that might be confounded by the appearance of additional peaks at longer mixing times.

**Figure 3.**
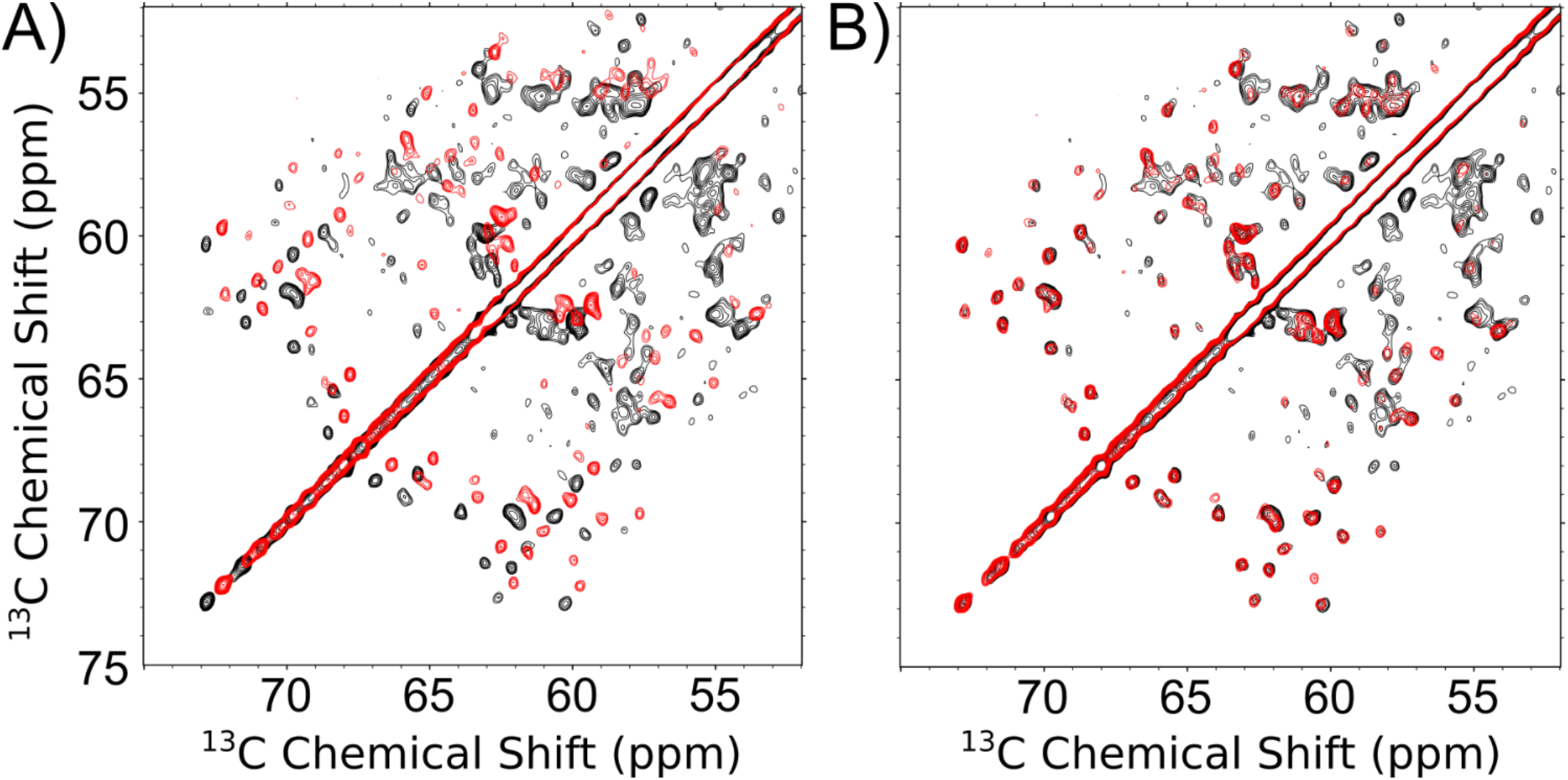
Alignment of spectra with different mixing times and instrument configurations of tryptophan synthase, a 144 kDa (72 kDa asymmetric unit) complex. (A) Overlaid unaligned tryptophan synthase ^13^C-^13^C spectra 300 ms CORD mixing time (black) with a correctly referenced tryptophan synthase ^13^C-^13^C spectra 100 ms CORD mixing (red). (B) Aligned spectra from (A) with a calculated shift of (0.580, 0.609) ppm. The cross-correlation score increases from 0.83 to 0.93.

**Figure 4.**
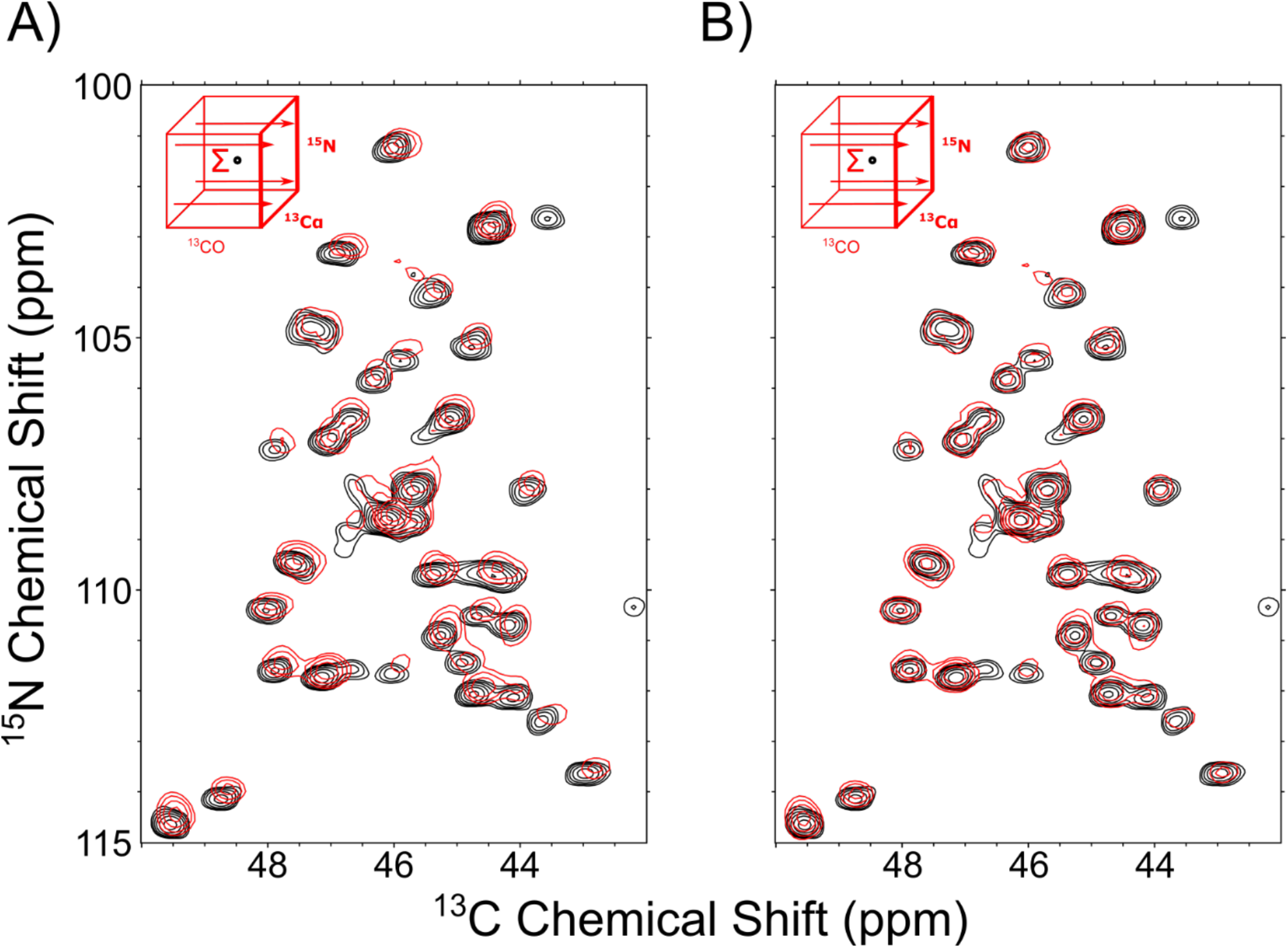
Alignment of 2D and 3D tryptophan synthase spectra with two common dimensions. Alignment of 3D NCAco sum projection with 2D NCA spectra dimensions. (A) unaligned NCAco projection (red) overlaid with NCA spectra (black). (B) Aligned NCAco projection and NCA spectra. Final shift: 0.100, 0.079 ppm. The cross-correlation score was 0.60 before alignment and 0.63 after.

Finally, we explored *ANSA*’s capability to address the challenging case of 3D spectral alignment. Correctly aligning 3D spectra with 2D spectra is essential for biomolecular NMR assignments but presents unique challenges due to the additional dimension and typically lower signal-to-noise ratios in 3D experiments. To address this, *ANSA* implements a projection-based approach, generating maximum-value projections from 3D datasets that can be aligned with corresponding 2D spectra sharing common dimensions. We tested this functionality using a tryptophan synthase NCACO 3D spectrum projected through the CO dimension (NCAco) for alignment with a 2D NCA spectrum. Despite both spectra being nominally “correctly referenced,” visual inspection revealed clear misalignment (Figure 3A)—a common occurrence even with careful experimental setup. *ANSA* detected a shift of (0.100, 0.079) ppm, producing well-aligned spectra (Figure 3B) and improving the cross-correlation score from 0.60 to 0.63. Notably, this 3D alignment example highlights *ANSA*’s ability to handle complex cases where different dimensions may require different referencing corrections, and the set of peaks observed in each spectrum is incomplete, as required for obtaining new assignments by identification of new peaks.

Notably, the LOW-BASHD decoupling scheme used during acquisition of the directly detected dimension of the 3D experiment introduced a shift offset distinct from that of the indirect dimensions – a subtlety that could be difficult to identify and correct through manual alignment. This offset reflects off-resonance effects inherent to the specific homonuclear decoupling pulses employed; while more complex pulse designs can minimize these effects, they persist in the simplest and most power-efficient implementations^8^.

*ANSA*’s resampling capabilities ensured proper alignment despite these complexities. Furthermore, we confirmed that *ANSA* provides consistent results even when spectra are processed with different spectral widths, demonstrating the algorithm’s robustness to common variations in spectral processing parameters. In all test cases, *ANSA* provided objective, reproducible alignment that eliminated the subjectivity inherent in manual methods. The algorithm’s performance remained consistent across varied experimental conditions, different types of spectroscopic experiments, and in the presence of noise and peak variations. These results demonstrate that *ANSA* offers a reliable solution to the persistent challenge of NMR spectral alignment, providing accuracy that meets or exceeds that of manual alignment while eliminating operator bias and improving workflow efficiency.

## CONCLUSIONS

We have developed Automated NMR Spectral Alignment (*ANSA*), a robust computational tool that successfully adapts cross-correlation function algorithms from cryoEM motion correction to address a persistent challenge in NMR spectroscopy. Through systematic testing across multiple scenarios, *ANSA* demonstrates significant advantages over traditional manual alignment methods. First, *ANSA* achieves perfect detection accuracy (0.1 ppm) in controlled test cases with artificially induced shifts, resulting in cross-correlation improvements from 0.33 to 1.00. More importantly, when applied to real experimental data, *ANSA* successfully aligned spectra collected months apart from different noise profiles, improving cross-correlation from 0.46 to 0.96. The algorithm also effectively handles spectra collected with different experimental parameters, such as varying mixing times, where peaks may differ substantially between datasets. Additionally, *ANSA* extends alignment capabilities to 3D datasets through strategic projections, enabling critical cross-validation between 2D and 3D experiments that form the foundation of modern biomolecular NMR analysis. Unlike manual alignment, which relies on subjective visual inspection and can be biased toward prominent peaks, *ANSA* provides objective, whole-spectrum alignment that considers all spectral features equally. This approach eliminates confirmation bias and ensures reproducibility across operators and facilities—a critical requirement for standardizing NMR data processing. The global cross-correlation approach proves particularly valuable for complex protein spectra containing thousands of peaks, where manual alignment becomes impractical. The implementation of *ANSA* as open-source software with an accessible interface ensures immediate integration into existing NMR processing workflows. This automation of a previously manual and subjective step represents an important advance toward more rigorous, reproducible NMR data analysis. Future developments could extend *ANSA*’s capabilities to directly align 3D volumes, incorporate weighted alignment approaches for spectra with dramatically different peak distributions, and enable batch processing for facility-wide referencing standardization. By bridging methodologies between cryoEM and NMR, *ANSA* demonstrates the value of cross-disciplinary algorithm adaptation in structural biology. As increasingly complex biomolecular systems are studied using integrative structural approaches, tools like *ANSA* that ensure precise data alignment become essential for extracting maximum information from complementary techniques.

## ASSOCIATED CONTENT

### Supporting Information

There is no supporting information for this article.

## AUTHOR INFORMATION

### Author Contributions

The manuscript was written through the contributions of all authors. All authors have given approval to the final version of the manuscript.

### Notes

The authors declare no competing financial interest.

## ACKNOWLEDGMENT

This study made use of the National Magnetic Resonance Facility at Madison (NMRFAM), an NIH Biomedical Technology Development and Dissemination Center (P41GM136463). The 1.1 GHz NMR spectrometer was funded by the United States National Science Foundation (NSF) Mid-Scale Research Infrastructure Big Idea (1946970). Helium recovery equipment, computers, and infrastructure for data archive were funded by the University of Wis-consin-Madison, NIH (P41GM136463, R24GM141526), and NSF (1946970). L.J.M. was supported by the NIH (R01GM137008 and R35GM145369).

